# Mechanism of stepwise electron transfer in six-transmembrane epithelial antigen of the prostate (STEAP) 1 and 2

**DOI:** 10.1101/2021.12.23.474010

**Authors:** Kehan Chen, Lie Wang, Jiemin Shen, Ah-lim Tsai, Ming Zhou, Gang Wu

## Abstract

Six transmembrane epithelial antigen of the prostate (STEAP) is a family of membrane-embedded hemoproteins with four members, STEAP1-4, all of which have a transmembrane domain (TMD) that chelates a heme prosthetic group. STEAP2-4, but not STEAP1, have an intracellular oxidoreductase domain (OxRD) so that an electron transfer chain composed of NADPH, FAD, and heme is established to mediate electron transfer across cell membranes. However, it is not known whether STEAP1 can establish a physiologically relevant electron transfer chain. Here we show that reduced FAD binds to STEAP1 and enables reduction of the heme. We also show that a soluble cytochrome *b*_5_ reductase can dock on STEAP1 and serve as a surrogate OxRD to reduce the heme. These results provide the first evidence that STEAP1 can support a cross-membrane electron transfer chain. It is not clear whether FAD, which relays electrons from NADPH to heme and interacts with both OxRD and TMD, remains constantly bound to the STEAPs. We found that FAD reduced by STEAP2 can be utilized by STEAP1, supporting the hypothesis that FAD is diffusible rather than staying bound to STEAP2. We determined the structure of human STEAP2 in complex with NADP^+^ and FAD to an overall resolution of 3.2 Å by cryo-electron microscopy. The structure of STEAP2 shows that the two cofactors bind similarly to those in the STEAP4 structure and thus a diffusible FAD is likely a general feature of the electron transfer mechanism in the STEAPs. The structure of STEAP2 also shows that its extracellular regions are less structured than those of STEAP4 or STEAP1, and further experiments show that STEAP2 reduces Fe^3+^-NTA with a rate significantly slower than STEAP1. These results establish a solid foundation for understanding the function and mechanisms of STEAP family of proteins.

## Introduction

<u>S</u>ix transmembrane <u>e</u>pithelial <u>a</u>ntigen of the <u>p</u>rostate 1 (STEAP1) was first discovered owing to its high level of expression in prostate cancer cells.^1^ Discovery of STEAP2-4 soon followed, and further analyses show that STEAP2-4 have metal ion reductase activities.^2, 3^ STEAP3 was identified as a ferrireductase required for iron uptake in erythroid cells.^3, 4^ STEAP2-4 are also found overexpressed in many types of cancer cells, suggesting their involvement in cancer initiation or progression.^1, 5, 6^

Each STEAP protein has a transmembrane domain (TMD) which consists of six transmembrane helices (TM), and STEAP2-4, but not STEAP1, also have a N-terminal intracellular oxidoreductase domain (OxRD).^7, 8^ TMD of the STEAP family of proteins are homologous to membrane-embedded reductases including mammalian NADPH oxidases (NOX) and dual oxidases (DUOX), and bacterial and yeast ferric reductases (FRE).^9^ The structures of NOX and DUOX show that their TMD binds two heme prosthetic groups, each ligated to a pair of conserved histidine residues from TMs 3 and 5; one heme is close to the intracellular side and the other close to the extracellular side of the TMD.^10–14^ In NOX or DUOX, the OxRD binds both NADPH and FAD, and the cross-membrane electron transfer chain starts with hydride transfer from NADPH to FAD, and then sequentially to the intracellular and extracellular hemes, and finally to the substrate. The structure of STEAP4, on the other hand, shows that only one heme is present in the TMD and it corresponds to the extracellular heme in NOX or DUOX.^15^ In STEAP4, the FAD straddles OxRD and TMD with the isoalloxazine ring of FAD binding at the equivalent position of the intracellular heme in NOX and DUOX and the nucleotide moiety of FAD binding to the OxRD (Fig. S1). While this configuration allows electron transfer between FAD and heme, the isoalloxazine ring is too far away from the nicotinamide ring of NADPH to receive hydride. Thus, the isoalloxazine ring of FAD must dissociate from the TMD and move closer to the nicotinamide ring of NADPH for hydride transfer. Although the isoalloxazine ring of FAD must assume different conformations during the redox cycles in STEAPs, its adenosine moiety could stay bound to the OxRD. However, here we show evidence that FAD does not stay tightly bound to the STEAP protein and can become diffusible after its reduction.

STEAP1 has been considered not be able to establish an electron transfer chain due to its lack of an OxRD, however, recent structures of STEAP1 and STEAP4 offer some hints otherwise.^15, 16^ In the STEAP4 structure, the isoalloxazine ring of FAD is coordinated by residues in TMD that are conserved in STEAP1. The structure of STEAP1 aligns well with that the TMD of STEAP4, and although FAD is not present in the STEAP1 structure,^16^ we showed previously that purified STEAP1 contains residual FAD, and that FAD binds to STEAP1 with K_D_ of ∼32 µM.^17^ Building upon these knowledge, we focused on identifying conditions that allow formation of a transmembrane electron transfer chain in STEAP1. We also determined the structure of human STEAP2, which shows that it has a more flexible substrate binding site compared to STEAP1 and STEAP4.

## Results

### Reduction of STEAP1 by reduced FAD

We first examined if the heme on STEAP1 can be reduced by FAD. We found that reduced FAD (as FADH^-^) readily reduces STEAP1, shown by the fast decrease in the Soret absorbance of ferric heme (A_413_) and the concomitant increase in the Soret absorbance of ferrous heme (A_427_) (Fig. 1A). The time course of A_427_ increase is biphasic. The fast phase has a rate constant of 7.7 s^−1^ and accounts for 60% of the total change at A_427_, and the slow phase, 0.67 s^−1^ and 40% (Fig. 1B). The rate constants of both phases exhibit dependence on [FADH^−^]. The *v*_max_ and K_M_ are estimated 12 s^−1^ and 4.7 μM for the fast phase, and the parameters are estimated 0.9 s^−1^ and 2.7 μM for the slow phase (Fig. 1B, inset).

**Figure 1.**
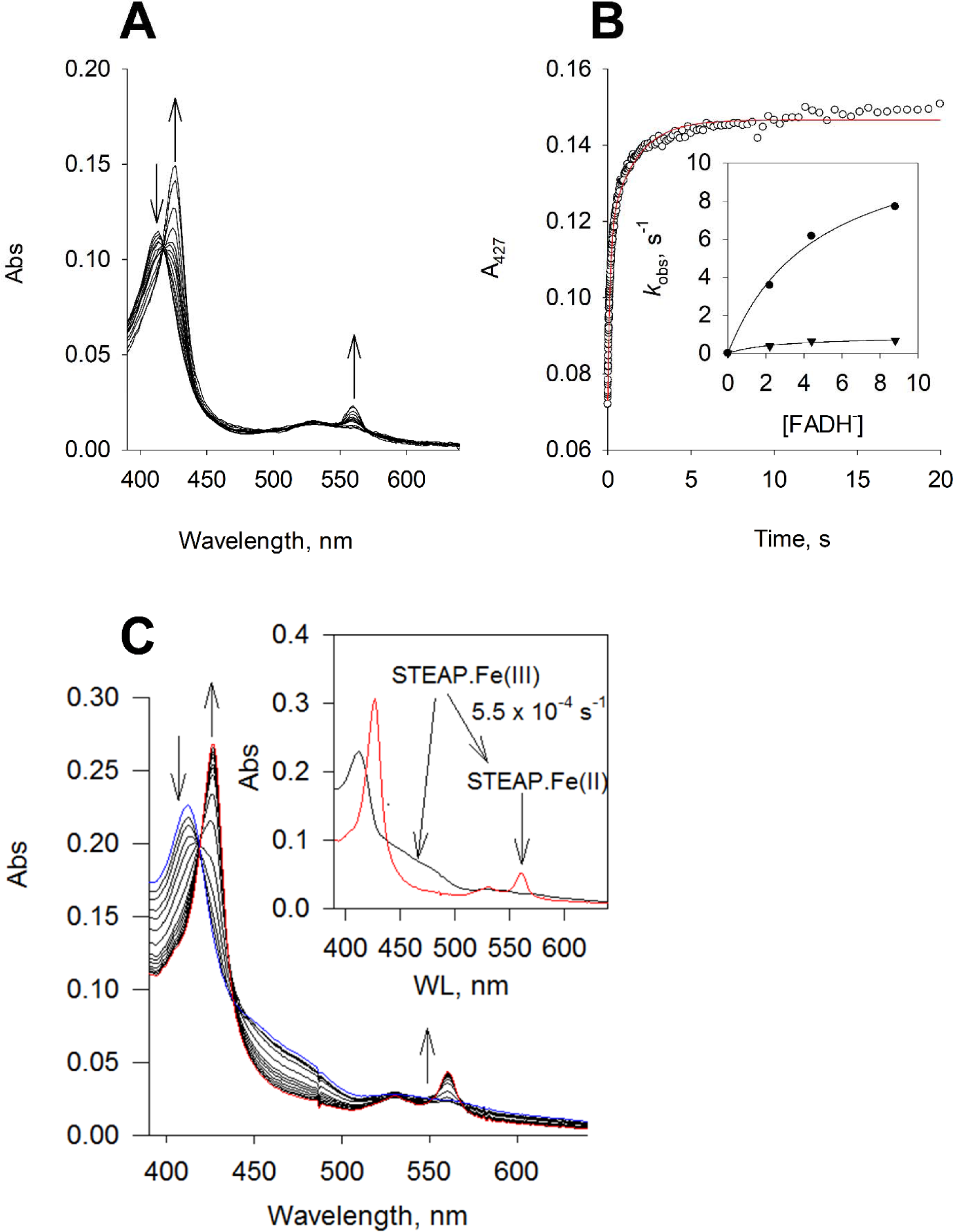
Reduction of the heme on STEAP1. (A) Rapid-scan reaction of 1.1 μM STEAP1 with 4.5 μM reduced FAD (FADH^−^); the spectral change was monitored for 20 s. (B) The time course of A_427_ (black), the Soret absorbance of ferrous heme, was extracted from the rapid-scan data. Red: biphasic exponential fit with rate constants *k*_obs_ of 7.7 (± 0.30) and 0.67 (± 0.034) s^−1^, respectively (n = 3). The percentage of each phase is 60% and 40%, respectively. Inset, the dependence of rate constants on [FADH^−^]. Dot, the fast phase; triangle, the slow phase. Lines, fit with equation *k*_obs_ = *v*_max_ * [FADH^−^]/(K_M_ + [FADH^−^]). (C) The spectral changes in the reaction of a mixture of 1.1 μM STEAP2 and 0.9 μM STEAP1 (plus 2.2 μM FAD) with 60 μM NADPH; the spectral change was monitored for 1 hr. The direction of the spectral changes is indicated by the arrows. Blue, the spectrum captured at the start of the reaction; red, the spectrum after 1 hour reaction. Inset, the resolved spectral species by deconvolution and the conversion rate constant. Black, ferric STEAP and red, ferrous STEAP.

In STEAP4 structure, a phenylalanine side chain (Phe359) is positioned between the isoalloxazine ring of FAD and the heme and is thought to mediate electron transfer from FAD to heme.^15^ In STEAP1-3, the equivalent residue is a leucine, Leu230 in STEAP1 (Fig. S2A). To examine if Leu230 is involved in electron transfer, we expressed and purified the L230G STEAP1 mutant. The UV-Vis spectrum of L230G STEAP1 is identical to that of the wild-type (WT) STEAP1 (Fig. S2B), indicating that the mutation does not perturb heme binding. When FADH^−^ was added to L230G STEAP1, biphasic reduction of heme was observed and the *k*_obs_ ^’^s showed dependence on [FADH^−^]. The *v*_max_ and K_M_ are estimated 2 s^−1^ and 3.6 μM for the fast phase, and the parameters are estimated 0.16 s^−1^ and 1.1 μM for the slow phase (Fig. S2C). These results indicate that the heme in L230G STEAP1 is reduced by FADH^−^ more than 5 times slower than in WT STEAP1, suggesting that the side chain of Leu230 is involved in mediating electron transfer between FAD and heme in STEAPs.

### Reduction of STEAP1 by cytochrome b_5_ reductase

We proceeded to test whether the heme on STEAP1 can be reduced by cytochrome *b*_5_ reductase (*b*_5_R). *b*_5_R catalyzes the reduction of a tightly bound FAD by NADH and is known to reduce the heme on cytochrome *b*_5_ (C*b*_5_).^18, 19^ Under anaerobic conditions, STEAP1 does not react with NADH but in the presence of both *b*_5_R and NADH, the A_427_ absorbance increases indicating reduction of the heme on STEAP1 (Fig. S3A). Using rapid-scan stopped-flow method, we captured the kinetics of the reactions when STEAP1 pre-incubated with *b*_5_R was mixed with NADH (Fig. 2A). The reduction of STEAP1 is clearly indicated by the shift of Soret peak (A_413_ to A_427_) and the split and increase of the α and β bands at 560 nm and 530 nm, respectively (Fig. 2A). Three spectral species, *A*, *B*, and *C* are resolved with rate constants of 177.9 (± 35.3) s^−1^ (*A* to *B*) and 0.13 (± 0.006) s^−1^ (*B* to *C*), respectively (Fig. 2A, inset). Spectral species *A* corresponds to ferric STEAP1 plus oxidized *b*_5_R while species *C* is ferrous STEAP1 with fully reduced *b*_5_R. A spectral intermediate species *B* can be identified with decreased absorbance in 420 – 500 nm range but little or no change in the Soret range when compared to *A* (Fig. 2A, inset), indicating that the intermediate *B* contains partially reduced *b*_5_R and a ferric heme. The resolution of intermediate *B* is due to fast FAD reduction in *b*_5_R by NADH without significant STEAP1 reduction. The *B* to *C* conversion reflects electron transfer from reduced *b*_5_R to STEAP1 (Fig. 2A), indicating that STEAP1 can form an electron transfer chain with *b*_5_R.

**Figure 2.**
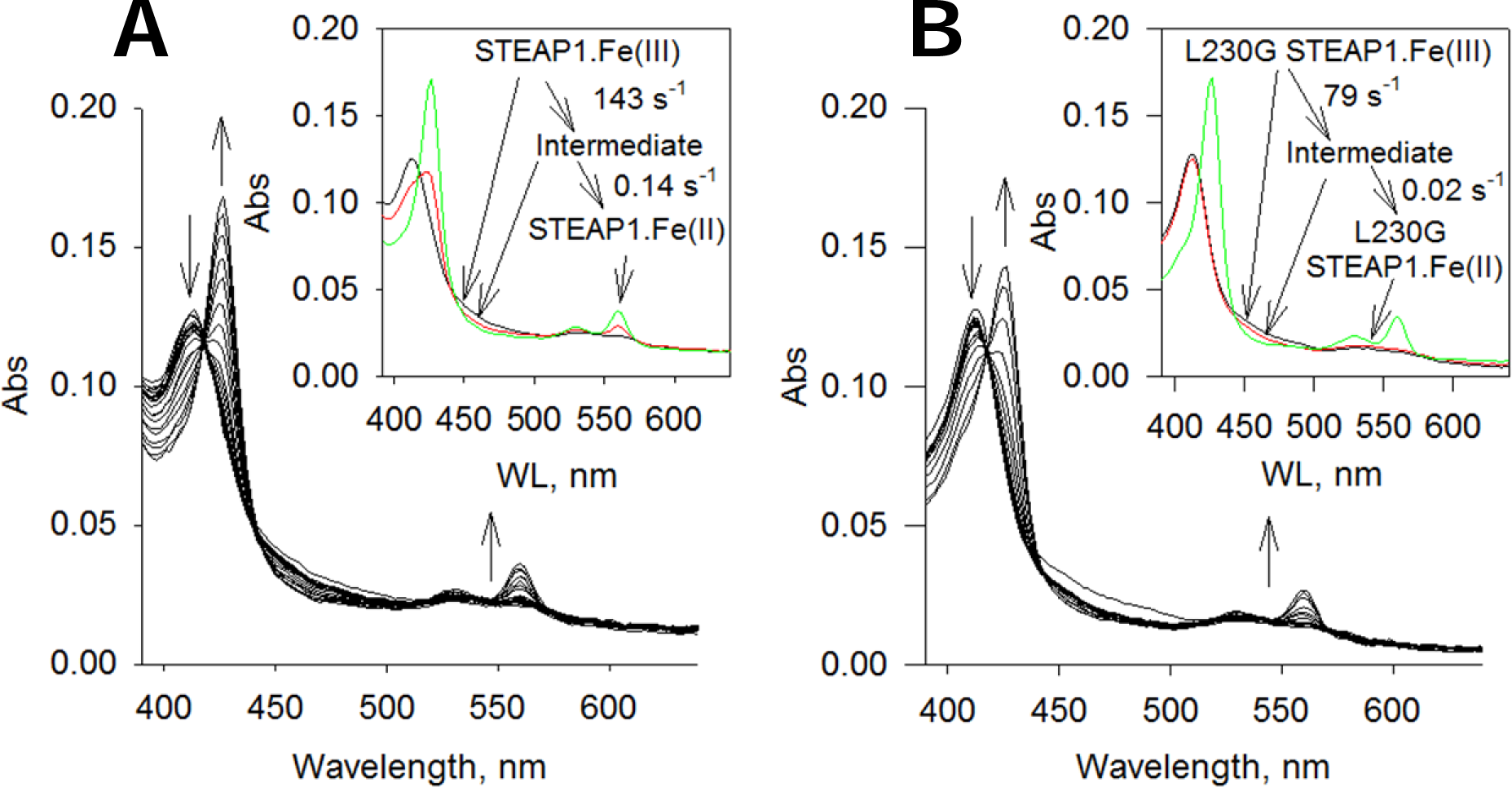
Reduction of STEAP1 by cytochrome *b*_5_ reductase. (A) The rapid-scan reaction of 1.5 μM STEAP1 and 1.5 μM cytochrome *b*_5_ reductase (*b*_5_R) with 10 μM NADH; the spectral change was monitored for 20 s. The arrows indicate the direction of the spectral change. Inset: the resolved spectral species and the conversion rate constants. Black, ferric STEAP1with *b*_5_R, red, a spectral intermediate, and green, ferrous STEAP1with fully reduced *b*_5_R. (B) L230G STEAP1 and *b*_5_R were reacted with 10 μM NADH; the spectral change was monitored for 50 s. The direction of spectral change is indicated by the arrows. Inset, the resolved spectral species by deconvolution and the rate constants. Inset: black, ferric L230G STEAP1 with *b*_5_R, red, a spectral intermediate, and green, ferrous L230G STEAP1 with fully reduced *b*_5_R.

We next examined whether *b*_5_R forms a complex with STEAP1. Using bio-layer interferometry assay (BLI), we measured the affinity between *b*_5_R and STEAP1 and found that *b*_5_R binds STEAP1 with a K_D_ of ∼5.9 µM (Fig. S4). We noticed that the fits to the association and dissociation profiles in BLI are rather poor at high concentrations of *b*_5_R and this is likely due to non-specific interactions between STEAP1 with *b*_5_R. Nonetheless, our BLI data demonstrates that *b*_5_R can dock onto STEAP1 to establish an electron transfer chain.

We also examined reduction of L230G STEAP1 by NADH and *b*_5_R (Fig. 2B). Three spectral species, *A*, *B*, and *C* are resolved, and the rate constants are 78.8 (± 22.6) s^−1^ (*A* to *B*) and 0.02 (± 0.01) s^−1^ (*B* to *C*) (Fig. 2B, inset). Species *A* corresponds to ferric L230G STEAP1 with oxidized *b*_5_R while species *C* represents ferrous L230G STEAP1 with fully reduced *b*_5_R (Fig. 2B, inset). As in WT STEAP1, the fast reduction of *b*_5_R by NADH leads to the resolution of spectral intermediate *B*, which has decreased absorbance between 420 – 500 nm but approximately the same Soret absorbance compared to *A* (Fig. 2B, inset). On the other hand, the rate constant from *B* to *C*, which is electron transfer rate from reduced *b*_5_R to the heme, is significantly slower in L230G STEAP1 than in WT STEAP1 (Fig. 2B, inset), suggesting that Leu230 is involved in the electron transfer from *b*_5_R to STEAP1.

### Purification and characterization of STEAP2

While structures full-length STEAP1 and STEAP4 have been determined by cryo-electron microscopy,^15, 16^ those of full-length STEAP2 and STEAP3 remain unresolved. We expressed and purified human STEAP2, which elutes as a single peak in size-exclusion chromatography and the elution volume is consistent with STEAP2 being a homotrimer (Fig. S5A). A prominent heme absorption peak is present in the purified STEAP2, and the heme content typically ranges from 70 – 90%. FAD is also detected in the purified STEAP2; however, at a level typically lower than 20% based on the fluorescence of FAD released from denatured STEAP2. No NADP(H) is detected in the purified STEAP2, suggesting that its association with STEAP2 is more transient than either heme or FAD.

The UV-Vis spectrum of STEAP2 heme are identical to that of STEAP1 heme. Ferric STEAP2 shows a Soret band at 413 nm and a broad Q band centered around 550 nm while the Soret band in ferrous STEAP2 shifts to 427 nm and the α and β bands are resolved at 560 and 532 nm, respectively (Fig. S5B). We further characterized the heme using magnetic circular dichroism (MCD) spectroscopy. The MCD spectrum of ferric STEAP2 shows strong Soret signals between 404 nm to 419 nm and no high-spin charge-transfer signal at wavelength above 600 nm while ferrous STEAP2 shows a much weaker Soret band but a very strong α band from 554 nm to 562 nm (Fig. S5C), consistent with the intense A-term Faraday effect of a typical low-spin *b*-type heme. Combined, the spectroscopic data indicates a *bis*-imidazole ligated low-spin heme in both redox states of STEAP2, consistent with a role in mediating electron transfer.

We monitored the spectral changes in STEAP2 in the reaction with NADPH. STEAP2 was pre-incubated with equal molar amount of FAD and reacted anaerobically with NADPH (Fig. S6A). In this reaction, the Soret absorbance of heme shifts from that of ferric state (A_413_) to that of ferrous heme (A_427_) while the α and β absorptions split into well resolved peaks at 560 nm and 532 nm, respectively, indicating the reduction of heme (Fig. S6A). Two spectral species are resolved with a transition rate constant of 1.2 (± 0.2) × 10^−3^ s^−1^ (*A* to *B*). Spectral species *A* corresponds to ferric STEAP2 plus FAD and spectral species *B* represents ferrous STEAP2 with reduced FAD, respectively (Fig. S6A, inset). Under the current experimental conditions, no intermediate with reduced FAD and ferric heme is resolved, suggesting that the oxidation of FAD by heme is significantly faster than its reduction by NADPH.

### STEAP1 reduction by STEAP2

Following the characterization of STEAP2, we investigated whether the reduced FAD produced in STEAP2 is accessible to STEAP1. We prepared an anaerobic mixture of STEAP2 (pre-incubated with equal molar of FAD) and STEAP1 and then added NADPH (Fig. 1C). In this reaction, the Soret absorbance of heme shifts from 413 nm to 427 nm and finally a narrow peak is observed at 427 nm with no shoulder at 413 nm (Fig. 1C), indicating that the heme on both STEAP1 and STEAP2 is fully reduced. Two spectral species are resolved from the reaction of the STEAP mixture with NADPH, corresponding to ferric STEAP plus FAD and ferrous STEAP with reduced FAD, with a rate constant of 5.5 × 10^−4^ s^−1^ (Fig. 1C, inset). On the other hand, in the absence of STEAP2, only a small fraction of STEAP1 is reduced (Fig. S3B), which may come from non-enzymatic reduction of FAD by NADPH. Thus, the heme on STEAP1 can be reduced by NADPH in the presence of STEAP2, via the reduced FAD produced in the OxRD of STEAP2. This data suggests that the reduced FAD becomes diffusible and quickly finds its binding pocket in the TMD of STEAP1. Under the current experimental conditions, the rate-limiting step seems to be the production of reduced FAD in the OxRD of STEAP2. Moreover, we cannot resolve the difference between diffusion of reduced FAD from STEAP2 to STEAP1 versus repositioning of reduced FAD from the OxRD to TMD in STEAP2.

### Fe^3+^-NTA reduction by STEAP1 and STEAP2

Our ability to produce reduced STEAP1 and STEAP2 allows measurement of the electron transfer step between the heme and a substrate. When reduced STEAP1 was mixed with Fe^3+^-NTA, the A_427_ absorbance decreased, indicating that the ferrous heme is oxidized by Fe^3+^-NTA. The time courses of A_427_ show biphasic kinetics with a fast phase accounting for 85% of the total absorbance change and a slow phase accounting for 15% (Fig. 3A). Similar biphasic kinetics was previously observed in the reactions of ferrous STEAP1 with Fe^3+^-EDTA or Fe^3+^-citrate.^17^ We speculate that there are two STEAP1 populations in terms of the conformation of the substrate binding site or geometry of the heme, or both. The *k*_on_ and *k*_off_ rate constants of Fe^3+^-NTA are estimated based on the *k*_obs_ vs. [Fe^3+^-NTA] (Fig. 3B) and the K_D_ of Fe^3+^-NTA for the two populations of STEAP1 is calculated, 50 and 26.3 μM for the fast and slow phase, respectively (Table S1). The K_D_ values of Fe^3+^-NTA are similar to those of Fe^3+^-EDTA or Fe^3+^-citrate (Table S1).

**Figure 3.**
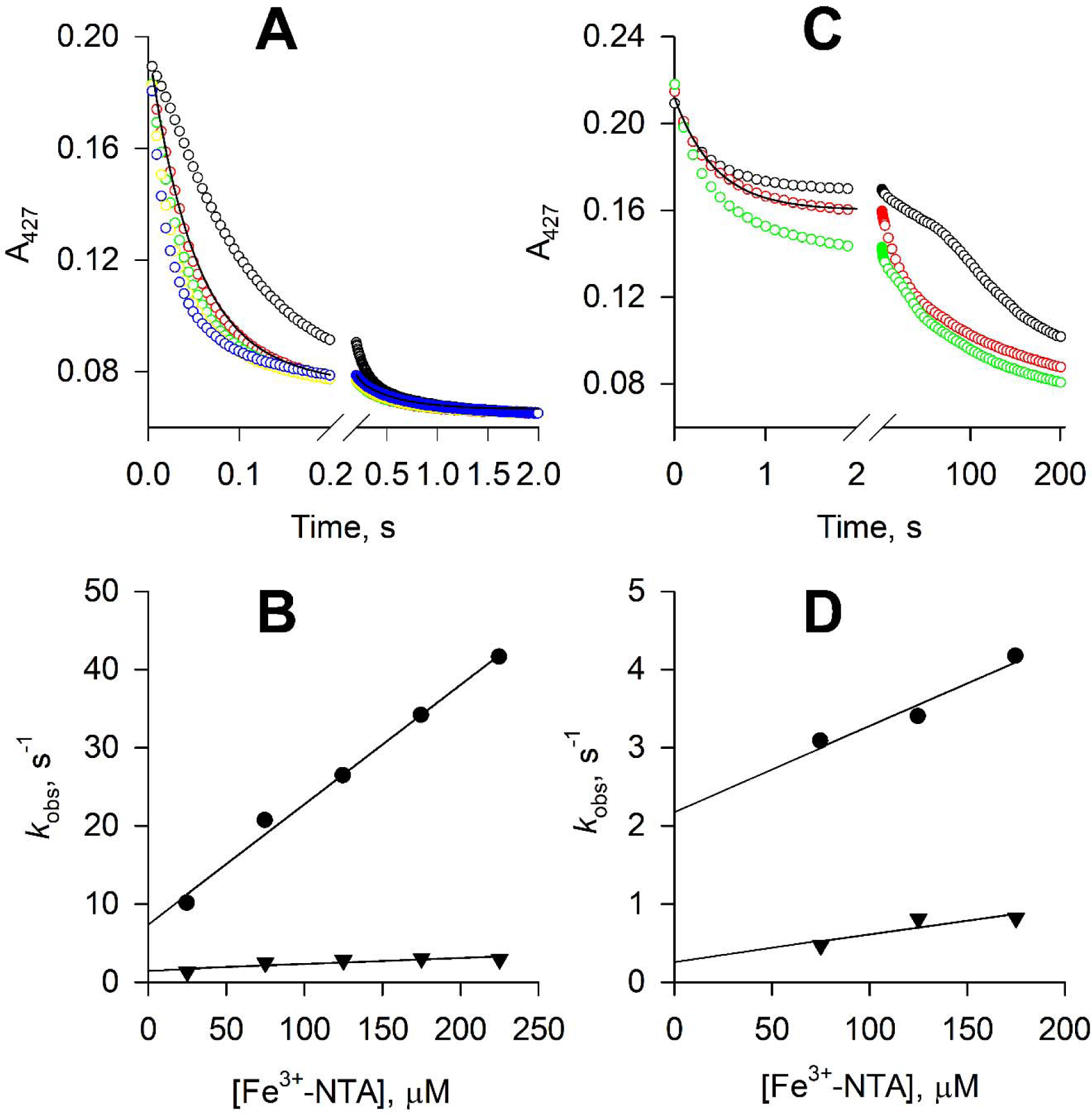
Reduction of Fe^3+^-NTA by ferrous STEAP1 and STEAP2. (A) The time courses of A_427_ in the reactions of 1.1 μM ferrous STEAP1 with 25 (black), 75 (red), 125 (green), 175 (yellow), and 175 μM Fe^3+^-NTA (blue). The rate constants, *k*_obs_, are estimated by biphasic exponential fit to the time courses. One of such fits is shown by the black line. (B) Dependence of the rate constants *k*_obs_ on [Fe^3+^-NTA]. Circles, *k*_obs_ of the fast phase of the A_427_ time courses; triangles, *k*_obs_ of the slow phase. (C) The time courses of A_427_ in the reactions of 1.1 μM ferrous STEAP2 with 75 (black), 125 (red), and 175 μM Fe^3+^-NTA (green). The time courses in the initial 2 s of the reactions are fitted with a biphasic exponential function, and one such fit is shown by the black line. (D) The rate constants estimated for the initial 2 s, *k*_obs_, are plotted versus [Fe^3+^-NTA]. Circles, *k*_obs_ of the fast phase of the A_427_ time courses; triangles, *k*_obs_ of the slow phase. At reaction time longer than 2 s, the time courses in (C) show more complicated kinetics and no clear dependence on [Fe^3+^-NTA].

The oxidation of ferrous STEAP2 by Fe^3+^-NTA is significantly slower than that of STEAP1 and the time courses of A_427_ show more than two phases (Fig. 3C), suggesting more heterogeneity in the substrate binding site or heme geometry. The ΔA_727_ in the initial 2 seconds accounts for about half of the total A_427_ decrease and the time courses in the initial 2 seconds can be fitted with a biphasic exponential decay function (Fig. 3C). The rate constants *k*_obs_ of the two phases show weak dependence on [Fe^3+^-NTA] (Fig. 3D). The *k*_on_ and *k*_off_ rate constants of Fe^3+^-NTA binding are estimated based on the *k*_obs_ vs. [Fe^3+^-NTA] plot (Fig. 3D) and the K_D_’s of Fe^3+^-NTA are calculated, 200 and 85.7 μM for the fast and slow phases, respectively (Table S1). After the initial 2 s, the time courses of A_427_ become slower with varying complicated shapes and have no clear dependence on [Fe^3+^-NTA] (Fig. 3D).

### Cryo-EM structure of STEAP2

We determined the structure of STEAP2 in the presence of NADP^+^ and FAD using cryo-EM. The cryo-EM data collection, refinement, and validation statistics are summarized in Table S2. The quality of density map is sufficient to build all the major structural elements of STEAP2 *de novo* with an overall resolution of 3.2 Å (Figs. S7 and S8). The N-terminal residues 1 – 27, C-terminal residues 470 – 490, and residues 332 – 353 (loop between TM3 and 4) are not resolved in our EM map, likely due to high degree of flexibility (Fig. 4D). Similar to the structure of STEAP4, STEAP2 has a domain-swapped homotrimer structure, where the OxRD of one protomer interacts with the TMD of a neighboring protomer (Fig. 4A and 4B). Overall, the structure of STEAP2 is very similar to that of STEAP4 with a root mean squared distance (RMSD) of 0.8 Å (Cα).^15^

**Figure 4.**
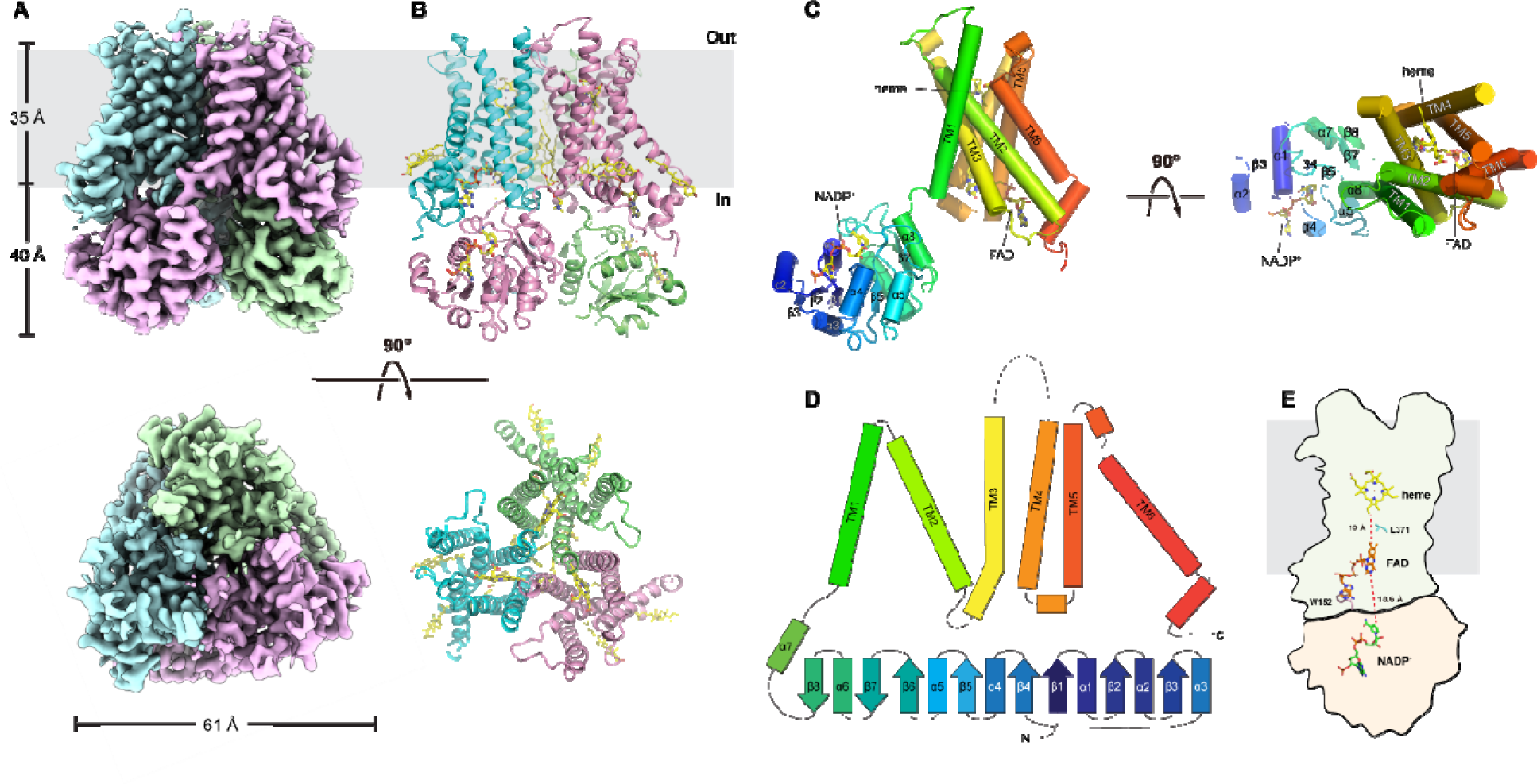
Cryo-EM structure of STEAP2. The sharpened density map (A) and cartoon presentation (B) for STEAP2 homotrimer. Top, the side view of STEAP2 homotrimer, and the grey bar represents the membrane; “in”, the intracellular side and “out’, the extracellular side. Bottom, the top view of STEAP2 homotrimer from the extracellular side. (C) The structure of one STEAP2 protomer (cartoon) with the prosthetic group heme and the cofactors FAD and NADP^+^ (sticks). Left, side view and right, top view from the extracellular side. (D) The topographic representation of the secondary structural elements. The α helices and β strands are represented by bars and arrows respectively. Dashed lines represent the unresolved segments. (E) The schematic representation of the spatial relationship of NADP^+^, FAD, and heme, shown as sticks. Trp152 and Leu371 are also shown as sticks. TMD is represented as the outline with grey shade and the OxRD with pink shade.

Heme, FAD, and NADP^+^ are unambiguously resolved in the density map (Figs. 4C and S7). The FAD molecule adopts an extended conformation as observed in STEAP4.^15^ The isoalloxazine ring buries deep in the TMD while the adenine ring of FAD forms a stacking interaction with Trp152 from the OxRD. The ribityl and pyrophosphate in FAD also interacts with residues on TMD. The isoalloxazine ring is ∼10 Å away (edge-to-edge) from the heme and the side chain of Leu371 protrudes approximately midway in between (Fig. 4E). Like L230 in STEAP1 (Fig. S2A), Leu371 may mediate electron transfer in STEAP2. The distance of the isoalloxazine ring of FAD to the nearest nicotinamide ring of NADP^+^ is ∼19 Å (Fig. 4E), which is too long for direct hydride transfer. We also found densities that likely correspond to phospholipid and cholesterol molecules. A 1-palmitroyl-2-oleoyl-glycero-3-phosphocholine molecule was built between the TMDs of two neighboring protomers, and two cholesterol molecules were built on the periphery of each TMD (Fig. S7). These tightly bound lipid molecules may have relevant structural and functional roles in STEAP2.

### Reduction of STEAP2 by reduced FAD

We measured reduction STEAP2 by reduced FAD in the absence of NADPH. Reacting with 4.5 μM reduced FAD (FADH^−^), ferric STEAP2 is reduced in biphasic kinetics with rates of 2.9 (± 0.80) s^−1^ (16%) and 6.9 (± 0.33) × 10^−2^ s^−1^ (84%), respectively (Fig. S6B). The reduction of STEAP2 by reduced FAD is significantly slower than that of STEAP1, and we attribute this to the presence of OxRD in STEAP2, which binds to the adenosine moiety of FAD but obstruct entrance of the isoalloxazine ring of reduced FAD into the TMD.

## Discussion

In this study, we demonstrate that STEAP1 is reduced by reduced FAD, either supplied directly or produced in the OxRD of STEAP2. We also show that *b*_5_R can reduce STEAP1 in the presence of NADH. Thus, STEAP1 may partner with various flavin-dependent reductases to establish an electron transfer chain from either NADH or NADPH to the extracellular side. These discoveries will facilitate our understanding of the physiological functions of STEAP1.

Purified STEAP2 has low levels of bound FAD, indicating that the FAD cofactor is not tightly bound as observed in other flavin reductases such as *b*_5_R or P450 reductase. In the structure of STEAP2, we captured the bound FAD in a conformation suitable for transferring electrons to heme, and the structure is similar to that of STEAP4.^15^ However, the structure of STEAP2 with the bound FAD in position to receive electrons from NADPH remains unresolved. Reduction of STEAP1 in the presence of STEAP2 provides the first evidence that the bound FAD may dissociate from STEAP2 after receiving electrons from NADPH. We propose that FAD first binds to OxRD of STEAP2 in a folded conformation with its isoalloxazine ring aligned with NADPH for hydride transfer. After receiving electrons from NADPH, reduced FAD may either dissociate from the OxRD and can be utilized by STEAP2 or STEAP1 nearby or stays partially bound to the OxRD and changes into an extended conformation with the isoalloxazine ring inserted deeply into the TMD (Scheme 1). A “diffusible” FAD or an FAD that switches between the folded and extended conformations will inevitably limit the rate of the overall electron transfer from NADPH to the extracellular side and is consistent with the kinetics data presented in Figs. 1C and S6A. This mechanism is different from the electron transfer chains found in the closely related NOX or DUOX enzymes, in which the FAD likely stays bound to the OxRD and thus suitable for rapid transfer of electrons from NADPH to the extracellular side. Moreover, the mechanism of regulating the activities of STEAPs *in vivo* must be very different from those in NOX or DUOX^20^ and warrants further investigation.

**Scheme 1.**
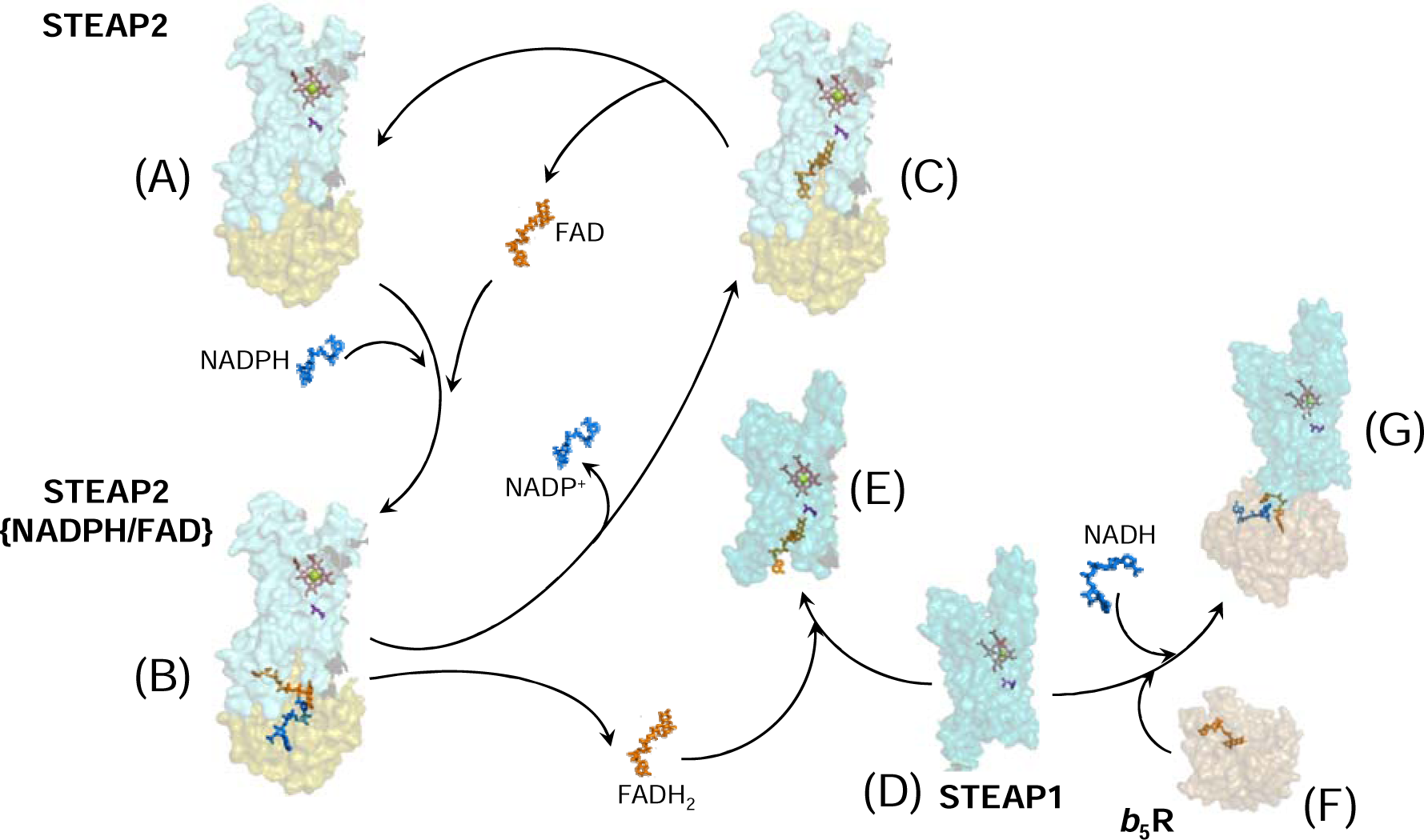
Electron transfer in STEAP1 and STEAP2. NADPH (blue) and FAD (orange) bind to the OxRD in STEAP2 (A, olive shade) with the nicotinamide ring of NADPH aligned with the isoalloxazine ring of FAD for hydride transfer (B). The reduced FAD adopts the extended conformation with its isoalloxazine ring bound deep in the TMD of STEAP2 (C, teal shade) or dissociates from the OxRD to bind STEAP1 (D ➔ E, cyan shade) and transfers electrons to heme (salmon). NADP(H) and FAD(H_2_) are co-factors that associate with and dissociate from the STEAP protein in each redox cycle while the heme, as a prosthetic group, stays bound to the protein. Cytochrome *b*_5_ reductase (F, sand shade) docks on STEAP1 from the intracellular side, forming a complex for electron transfer (G). The FAD-to-heme electron transfer in STEAP is likely mediated through a bulky sidechain (purple), Leu230 in STEAP1 and Leu371 in STEAP2, respectively.

We demonstrate that STEAP1 can establish electron transfer chain with *b*_5_R, which does not release FAD. We also show that *b*_5_R reduces STEAP1 from the intracellular side and it can form a complex with STEAP1. These results indicate that *b*_5_R can serve as a surrogate OxRD to complete the electron transfer chain. However, further analysis is required to establish whether STEAP1 pairs with *b*_5_R or other FAD reductase *in vivo*.

We are able to measure the rate of electron transfer from heme to a ferric substrate, and we found that STEAP2 reduces Fe^3+^-NTA significantly slower than STEAP1. The more complex time course of heme oxidation also suggests a high level of heterogeneity in either the substrate binding pocket or the heme geometry in STEAP2. In the cryo-EM structure of STEAP2, residues 332 – 353 between TM3 and 4 are unresolved, likely due to high flexibility in this region, while in the structures of STEAP1 and STEAP4, the corresponding residues form a well-defined extracellular loop adjacent to the putative substrate binding site.^15, 16^ These differences suggest that STEAP2 has different substrates from STEAP4.

## Materials and Methods

### Materials

4-(2-hydroxyethyl)-1-piperazineethanesulfonic acid (HEPES), 5-aminolevulinic acid (5-ALA), isopropyl β-D-1-thiogalactopyranoside (IPTG), phenylmethanesulfonyl fluoride (PMSF), Fe(NO_3_)_3_, hemin chloride, dithionite, 1-palmitroyl-2-oleoyl-glycero-3-phosphocholine (POPC), nitrilotriacetic acid (NTA), 1-ethyl-3-[3-dimethylaminopropyl]carbodiimide hydrochloride (EDC), and N-hydroxysulfosuccinimide (Sulfo-NHS) were from Sigma-Aldrich (St. Louis, MO). Lauryl maltose neopentyl glycol (LMNG) and glyco-diosgenin (GDN) were from Anatrace (Maumee, OH).

### Protein expression and purification

The human STEAP2 gene (NCBI accession number AAN04080.1) was codon optimized and cloned into a modified pFastBac Dual expression vector for production of baculovirus according to the Bac-to-Bac method (Thermo Fisher Scientific, Waltham, MA). P3 viruses were used to infect High Five (*Trichoplusia ni*) or Sf9 (*Spodoptera frugiperda*) insect cells at a density of ∼ 3 × 10^6^ cells mL^−1^ in the media including 0.5 mM 5-aminolevulinic acid, 10 µM FeCl_3_, and 5 µM hemin chloride. Infected cells were grown at 27 °C for 48 – 60 h before harvest. Cell membranes were prepared using a hypotonic/hypertonic wash protocol as previously described.^21^ Purified cell membrane pellets were then flash-frozen in liquid nitrogen for further use.

Purified membranes were thawed and homogenized in 20 mM HEPES, pH 7.5 buffer containing 150 mM NaCl, and then solubilized with 1.5% (w/v) LMNG at 4 °C for 2 h. After solubilization, cell debris was removed by ultracentrifugation (55,000 × g for 45 min at 4 °C), and hSTEAP2 was purified from the supernatant using cobalt-based Talon affinity resin (Clontech, Mountain View, CA). The C-terminal His_6_-tag was cleaved with tobacco etch virus (TEV) protease at room temperature for 30 min. The protein was concentrated to around 5 mg mL^−1^ using an Amicon spin concentrator with a 100 kDa cut-off (Millipore, Burlington, MA), and then loaded onto a SRT-3C SEC-300 size-exclusion column (Sepax Technologies, Newark, DE) equilibrated with 20 mM HEPES buffer containing 150 mM NaCl and 0.01% (w/v) LMNG. For the sample used in the cryo-EM structural studies, the size-exclusion column was equilibrated with 20 mM HEPES buffer containing 150 mM NaCl and 0.02% GDN.

Rabbit STEAP1 (NCBI accession number NP_001164745.1) was expressed and purified following the method published previously.^17^ The L230G STEAP1 mutation was introduced by the QuikChange method (Stratagene, CA) using the primers:

forward, 5’-CGTGGGACTGGCTATCGGCGCTTTGCTGGCTGTGAC-3’;

reverse, 5’-GTCACAGCCAGCAAAGCGCCGATAGCCAGTCCCACG-3’.

The cDNA of mouse cytochrome *b*_5_ reductase (*b*_5_R, UniProt Q3TDX8, soluble form, residues 24 – 301) was subcloned into a pET vector which appends a polyhistidine tag and a tobacco etch virus (TEV) protease site to the N-terminus of the overexpressed protein. The expression of *b*_5_R followed the previous protocol^22^ and the cell media was supplemented with 100 μM FAD.

### Electronic absorption and magnetic circular dichroism (MCD) spectroscopy

UV-Vis spectra of STEAP2 were recorded using a HP8453 diode-array spectrophotometer (Hewlett-Packard, Palo Alto, CA). The extinction coefficient of the heme was determined by pyridine hemochrome assay as published previously.^17^ MCD spectra of STEAP2 were recorded with a Jasco J-815 CD spectropolarimeter (Tokyo, Japan) equipped with an Olis permanent magnet (Bogart, GA). The parameters for MCD measurements are spectral bandwidth, 5 nm; time constant, 0.5 s; scan speed, 200 nm/min. Each MCD spectrum is an average of 12 repetitive scans and the signal intensity is expressed in units of M^−1^cm^−1^ tesla^−1^.

### Cryo-EM structure determination of STEAP2

Quantifoil R1.2/1.3 Cu grids were glow-discharged in air for 15 s at 10 mA in a plasma cleaner (PELCO EasiGlow, Ted Pella, Inc., Redding, CA). Glow-discharged grids were prepared using Thermo Fisher Vitrobot Mark IV. Concentrated hSTEAP2 in the presence of FAD and NADP^+^ (3.5 μl) was applied to each glow-discharged grid. After blotted with filter paper (Ted Pella, Inc.) for 4.0 s, the grids were plunged into liquid ethane cooled with liquid nitrogen. A total of 7509 micrograph stacks were collected using SerialEM^23, 24^ on a Titan Krios electron microscope (Thermo Fisher) at 300 kV with a Quantum energy filter (Gatan, Pleasanton, CA), at a nominal magnification of 105,000× and with defocus values of −2.5 μm to −0.8 μm. A K3 Summit direct electron detector (Gatan) was paired with the microscope. Each stack was collected in the super-resolution mode with an exposing time of 0.175 s per frame for a total of 50 frames. The dose was about 50 e^−^ per Å^2^ for each stack. The stacks were motion-corrected with MotionCor2^25^ and binned (2 × 2) so that the pixel size was 1.08 Å. Dose weighting^26^ was performed during motion correction, and the defocus values were estimated with Gctf.^27^

A total of 4,210,570 particles were automatically picked (RELION 3.1)^28–30^ from the motion-corrected images and imported into cryoSPARC^31^. After 2 rounds of two-dimensional (2D) classification, a total of 91 classes containing 1,031,895 particles were selected. A subset of 12 classes containing 117,053 particles were selected for *ab initio* three-dimensional (3D) reconstruction, producing one good class with recognizable structural features and three bad classes with no distinct structural features. Both the good and bad classes were used as references in the heterogeneous refinement (cryoSPARC) and yielded a good class at 4.10 Å from 305,849 particles. Non-uniform refinement (cryoSPARC) was then performed with C3 symmetry and an adaptive solvent mask, producing a map with an overall resolution of 3.16 Å. Resolutions were estimated using the gold-standard Fourier shell correlation with a 0.143 cut-off^32^ and high-resolution noise substitution.^33^ Local resolution was estimated using ResMap.^34^

The structural model of STEAP2 was built based on the cryo-EM structure of STEAP4 (PDB ID: 6HCY),^15^ and the side chains were adjusted based on the density map. Model building was conducted in Coot^35^. Structural refinements were carried out in PHENIX in real space with secondary structure and geometry restraints.^36^ The EMRinger Score was calculated as described previously.^37^

### STEAP reduction by NADPH

STEAP2, 2.3 μM, or a mixture of 1.1 μM STEAP2 and 0.9 μM STEAP1 was pre-incubated with 2.5 and 2.2 μM FAD in a tonometer, respectively. The solutions were made anaerobic by 5 anaerobic cycles, each with 30 s vacuum followed by argon sparging for 4.5 min. The stock solution of NADPH was made anaerobic by N_2_ sparging. Anaerobic NADPH solution was injected into the anaerobic STEAP solution using an airtight syringe and the spectral changes were monitored at room temperature using the HP 8453 spectrophotometer. The spectral changes were deconvoluted using the Pro-Kineticist program coming with the stopped-flow machine (see below).

### Stopped-flow experiments

To measure the electron transfer rate from ferrous STEAP to ferric nitrilotriacetic acid (Fe^3+^-NTA) substrate, anaerobic STEAP was first titrated to the ferrous state using dithionite and then reacted with Fe^3+^-NTA on an Applied Photophysics model SX-18MV stopped-flow machine (Leatherhead, UK), which was placed in a COY anaerobic chamber (Grass Lake, MI). The time course of A_427_, which reflects the oxidation of ferrous STEAP, was followed. Fe^3+^-NTA was prepared with ferric nitrate and NTA based on a ratio of [Fe^3+^]:[NTA] = 1:4. The rate constants of the redox reactions, *k*_obs_, were obtained by fitting the time courses using a monophasic or a multiphasic exponential function. The 2^nd^-order *k*_on_ rate constants of Fe^3+^-NTA to the STEAP protein were estimated from the slopes of the linear fits to the *k*_obs_ vs. [Fe^3+^-NTA] plots. On the other hand, the *k*_off_ rate constants were estimated based on the intercepts on ordinate of the linear fits.

The anaerobic protein mixture of STEAP1 and *b*_5_R was reacted with NADH and the spectral changes were monitored using a rapid-scan diode-array accessory with the stopped-flow machine. In the reaction of STEAPs with pre-reduced FAD, FAD in 20 mM HEPES, pH7.5 containing 150 mM NaCl and 0.1% LMNG was titrated anaerobically with dithionite. Most of the reduced FAD (FADH_2_) was likely in its ionized form FADH^−^ due its pKa = 6.7. Part of the reduced FAD was re-oxidized due to the very negative potential of the FAD/FADH^−^ pair, and the absorbance of FAD was subtracted as the background. The spectral changes were deconvoluted using the Pro-Kineticist program coming with the stopped-flow machine.

### Octet Bio-layer Interferometry

Bio-layer Interferometry (BLI) assays were performed at 30 °C under constant shaking at 1000 rpm using an Octet system (FortéBio, Fremont, CA). STEAP1 was immobilized on amine reactive second-generation (AR2G) biosensors (Sartorius, Göttingen, Germany). The biosensor tips were activated for 5 min in EDC and 10 mM Sulfo-NHS before being loaded with STEAP1 at a concentration of 1 µg/mL for 10 min. The tips were then quenched in 1 M ethanolamine (pH 8.5) for 5 min and equilibrated in 20 mM HEPES, pH7.5 containing 150 mM NaCl, 0.1% LMNG, and 0.1% BSA to reduce non-specific binding. The tips were then transferred into wells containing various concentration of *b*_5_R, 20, 10, 5, 2.5, 1.3, and 0.6 µM, for association and then back to the equilibration wells for dissociation. The binding curves were aligned and corrected with the channel with no analyst protein. The association and dissociation phases were fitted with a monophasic exponential function. The equilibrium responses (Req) in the association incubation were plotted against [*b*_5_R] and fitted with a dose-response function to calculate the dissociation constant K_D_ of STEAP/*b*_5_R complex.

## Data and Materials Availability

The EM data and fitted model of human STEAP2 are deposited in the Electron Microscopy Data Bank (EMD-25775) and the RCSB Protein Data Bank (7TAI). Original data is available upon request, please contact.

## Author Contribution Statement

GW and MZ initiated and designed the project. KC and LW led the effort of protein production and structure determination and with help from JS and GW. JS prepared protein mutant and measured protein-protein binding affinity. GW designed and conducted spectroscopic measurements. All authors analyzed and interpreted the data. GW and MZ wrote the manuscript with input from all authors.

