## Supplemental Materials for "Mechanism of stepwise electron transfer in six-transmembrane epithelial antigen of the prostate (STEAP) 1 and 2"

**Table S1 The rate constants of the reduction of ferric substrates by ferrous STEAP1 and 2.**

| | Substrates | First Phase | | Second Phase | |
|---|---|---|---|---|---|
| | | $k_{on}$, $M^{-1}s^{-1}$/$k_{off}$, $s^{-1}$ | $K_D$, μM* | $k_{on}$, $M^{-1}s^{-1}$/$k_{off}$, $s^{-1}$ | $K_D$, μM* |
| STEAP1† | $Fe^{3+}$-EDTA | $2.7 \times 10^5$ / 4.8 | 17.8 | $4.0 \times 10^4$ / 1.4 | 35 |
| STEAP1† | $Fe^{3+}$-citrate | $1.6 \times 10^5$ / 15.5 | 97 | - | - |
| STEAP1‡ | $Fe^{3+}$-NTA | $1.5 \times 10^5$ / 7.5 | 50 | $7.6 \times 10^3$ / 0.2 | 26.3 |
| STEAP2‡,¶ | $Fe^{3+}$-NTA | $1.1 \times 10^4$ / 2.2 | 200 | $3.5 \times 10^3$ / 0.3 | 85.7 |

*: $K_D = k_{off}/k_{on}$. †: from Kim, K. et al, *Biochemistry* (2016) **55**, 6673-6684. ‡: this study. ¶: the third phase is not included.

**Table S2 The data collection, refinement, and validation statistics of STEAP2 Cryo-EM.**

| | |
|---|---|
| **Data collection and processing** | |
| Magnification | FEI TITAN KRIOS |
| Voltage (kV) | 300 |
| Electron exposure ($e^{-}$/Å$^2$) | 50 |
| Defocus range (μm) | -0.8 to -2.5 |
| Pixel size (Å) | 1.08 |
| Symmetry imposed | C3 |
| Number of initial particle images | 4,210,570 |
| Number of final particle images | 117,053 |
| Map resolution (Å) | 3.16 |
| FSC threshold | 0.143 |
| Map resolution range (Å) | 3.2 |
| **Refinement** | |
| Initial model used | PDB 6hcy |
| Model resolution (Å) | 3.2 |
| FSC threshold | 0.5 |
| Map sharpening B factor (Å$^2$) | -100 |
| Model composition | |
| Non-hydrogen atoms | 11,109 |
| Protein residues | 1260 |
| Ligands | 18 |
| B factors (Å$^2$) | |
| Protein | 41.92 |
| Ligand | 32.76 |
| R.m.s. deviations | |
| Bond lengths (Å) | 0.003 |
| Bond angles (°) | 0.567 |
| Validation | |
| MolProbity score | 1.6 |
| Clashscore | 10.54 |
| Poor rotamers (%) | 0.09 |
| Ramachandran plot | |
| Favored (%) | 97.8 |
| Allowed (%) | 2.2 |
| Disallowed (%) | 0 |

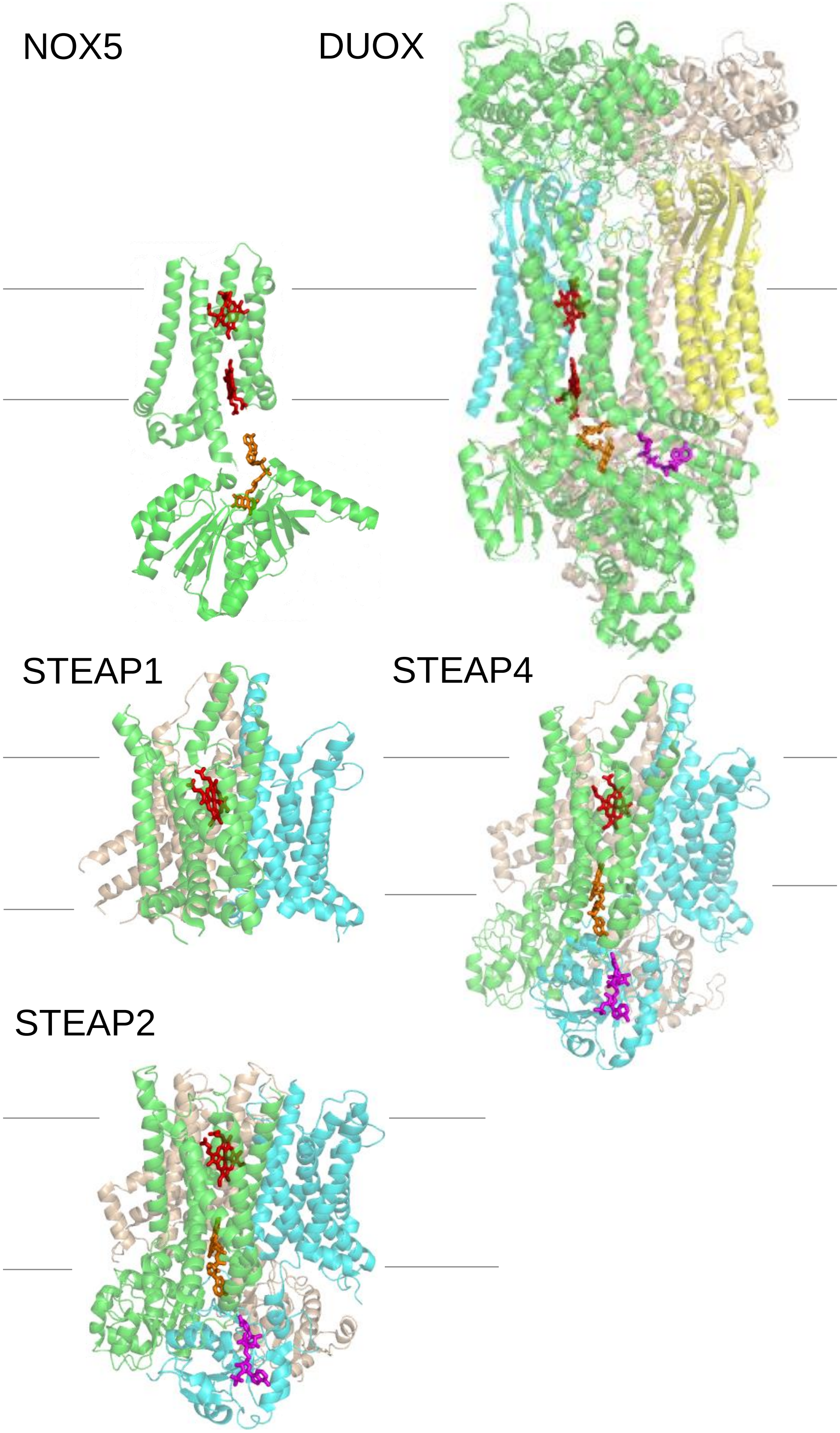
NOX5
DUOX
STEAP1
STEAP4
STEAP2

**Figure S1. The structures of NOX, DUOX, and STEAPs.** The crystal structures of the TMD and OxRD domains of NOX5 (from *Cylindrospermum stagnale*) are plotted together (PDB codes: TMD, 5o0t; OxRD, 5o0x). The cryo-EM structure of human DUOX1 (PDB code: 7d3f) shows dimeric oligomerization (green and cyan) complexed with DUOX auxiliary protein A1 (DUOXA1, wheat and yellow). The cryo-EM structures of STEAPs are homotrimers (green, cyan, and wheat. PDB codes: STEAP1, 6y9b; STEAP4, 6hcy; STEAP2, 7tai). In DUOX and STEAPs, only one set of cofactors and heme group are shown for clarity. Heme, red; FAD, orange; $NADP^+$, magenta. The lines represent cell membrane (top, extracellular and bottom, intracellular).

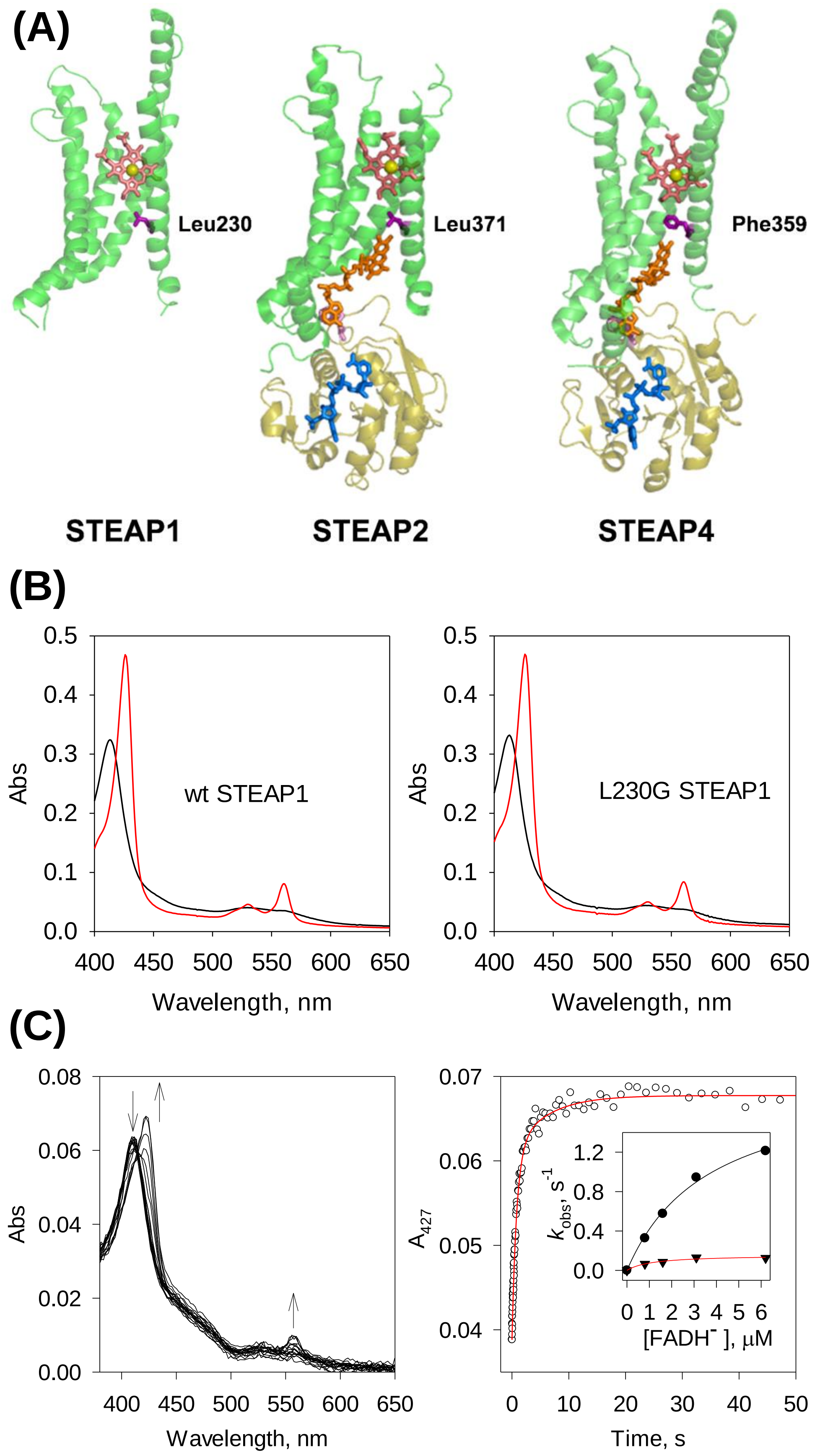
(A)
Leu230
Leu371
Phe359
STEAP1
STEAP2
STEAP4
(B)
Abs
wt STEAP1
L230G STEAP1
Wavelength, nm
(C)
Abs
Wavelength, nm
$A_{427}$
Time, s
$k_{obs}$, $s^{-1}$
[FADH⁻], μM

**Figure S2. Electron transfer in STEAPs may be mediated by a bulky sidechain.** (A) FAD (orange) binds both STEAP2 and STEAP4 in an extended conformation (PDB codes: STEAP2, 7tai; STEAP4, 6hcy). Its isoalloxazine ring protrudes deeply into the TMD domain (green) and the adenosine ring stacks with Trp152 or Trp140 (pink) in STEAP2 and STEAP4, respectively, from the OxRD of a neighboring protomer (olive). Leu371 or Phe359 (purple), in STEAP2 and STEAP4, respectively, lies halfway between the isoalloxazine ring and heme (salmon, iron atom in yellow). $NADP^+$ is represented in blue. In STEAP1 (PDB code: 6y9b), Leu230 (purple) is similarly positioned versus the heme. (B) The UV-Vis spectra of ferric (black) and ferrous (red) wt STEAP1 (left panel) and L230G STEAP1 (right panel). The ferrous proteins were prepared by photoreduction for 10 min. (C) Rapid-scan reaction of 0.4 μM L230G STEAP1 with 6.2 μM reduced FAD ($FADH^-$); the spectral change was monitored for 50 s (left panel). The arrows represent the direction of spectral changes. Right panel, the time course of $A_{427}$ extracted from the rapid-scan data. Red: biphasic exponential fit with rate constants $k_{obs}$ of 1.2 (± 0.13) and 9.6 × $10^{-3}$ (± 4.3 × $10^{-2}$) $s^{-1}$, respectively (n = 3). The percentage of each phase is 79% and 21%, respectively. Inset, the dependence of rate constants on [$FADH^-$]. Dot, the fast phase; triangle, the slow phase. Lines, fit with equation $k_{obs} = v_{max} * [FADH^-]/(K_M + [FADH^-])$.

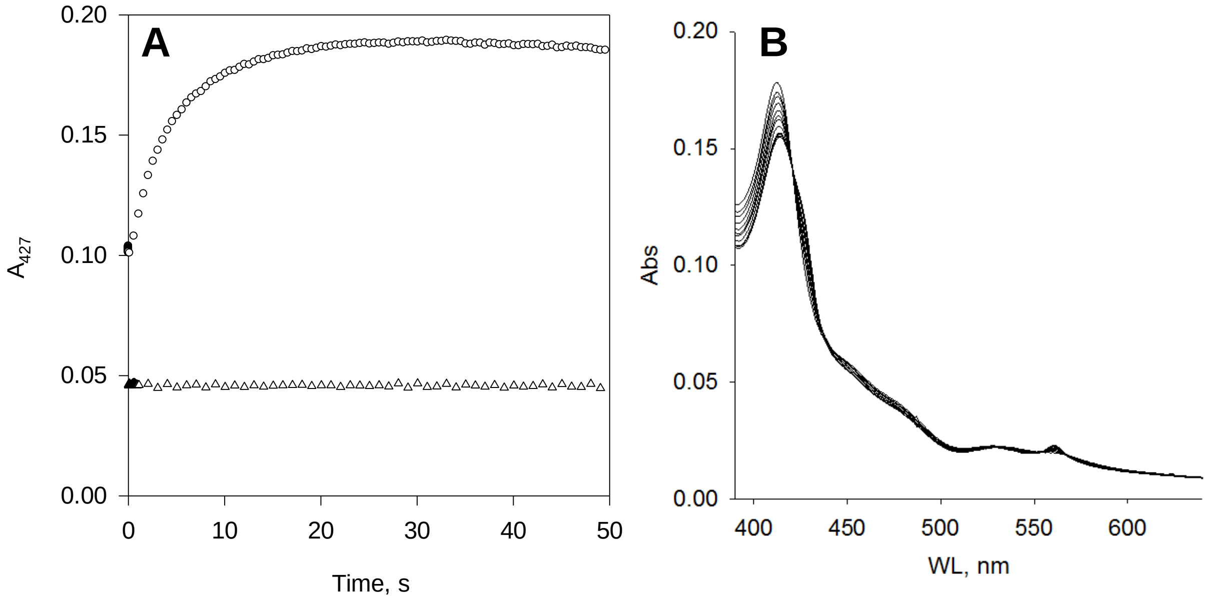


**Figure S3. STEAP1 in the presence of FAD and/or NAD(P)H.** (A) No reaction was observed when anaerobic 0.8 μM STEAP1 was mixed with 50 μM NADH (triangle). STEAP1 reduction was observed reacting anaerobic 1.7 μM STEAP1 pre-incubated with 1.3 μM $b_5$R with 1.3 μM NADH (circle). (B) Anaerobic STEAP1 plus FAD was incubated with NADPH for 1 hour.

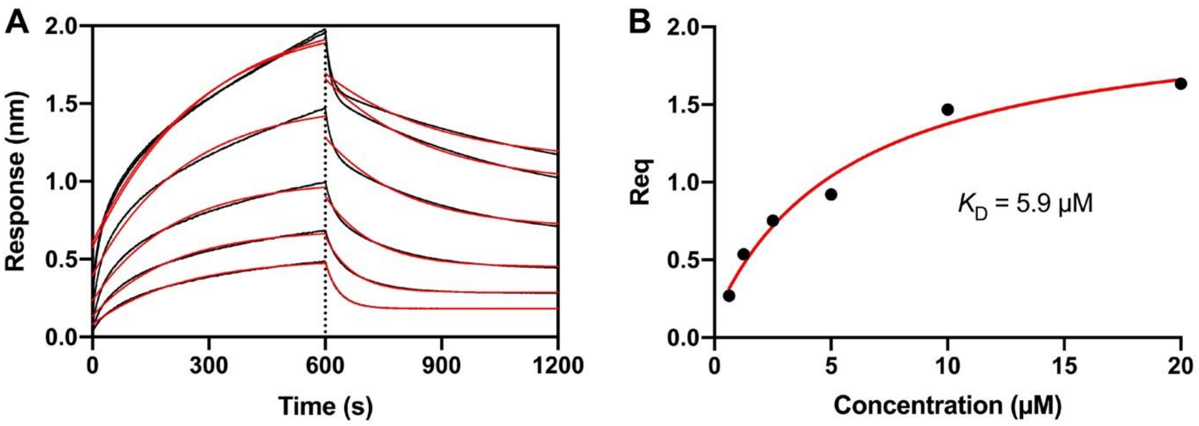


**Figure S4. Binding of *b*$_5$R to STEAP1.** (A) The Octet BLI sensorgram of various concentrations of *b*$_5$R binding to immobilized STEAP1. The original binding traces (black) were fitted with mono-exponential function (red). (B) Dose-response curve of the equilibrium response (Req) during association versus the concentration of *b*$_5$R. The dissociation constant $K_D$ was calculated from the fit (red line) to data (black dot). The $K_D$ is 5.9 μM (2.4 – 15.5 with 95% confidence interval).

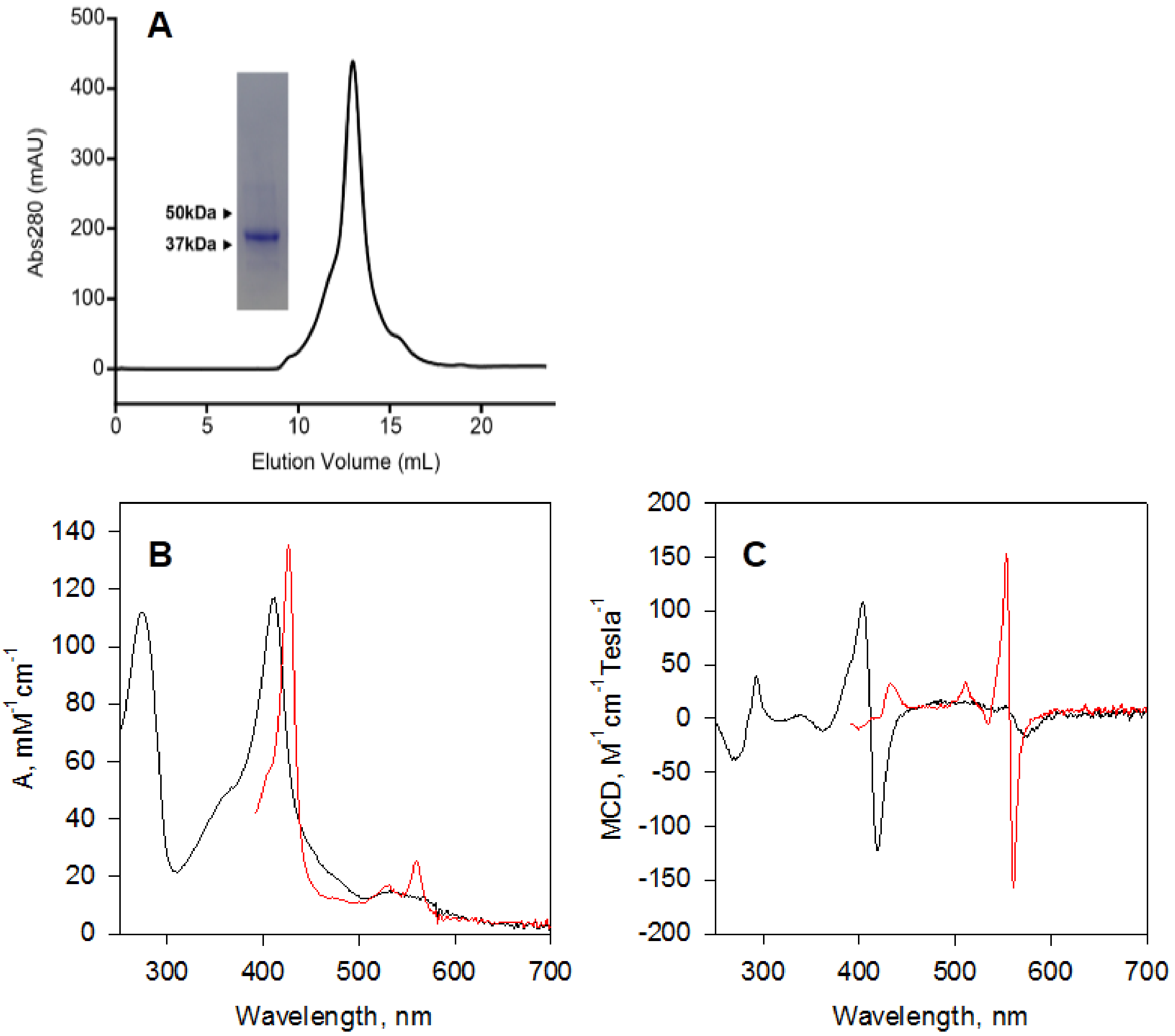


**Figure S5. Characterization of purified hSTEAP2.** (A) SDS-PAGE analysis and size exclusion chromatography of purified hSTEAP2. The UV-Vis (B) and MCD (C) spectra of hSTEAP2 are plotted with absorbance in $mM^{-1}cm^{-1}$ and MCD in $M^{-1}cm^{-1}Tesla^{-1}$, respectively. STEAP2 was purified in the ferric state (black) and the ferrous STEAP2(red) was prepared by dithionite reduction.

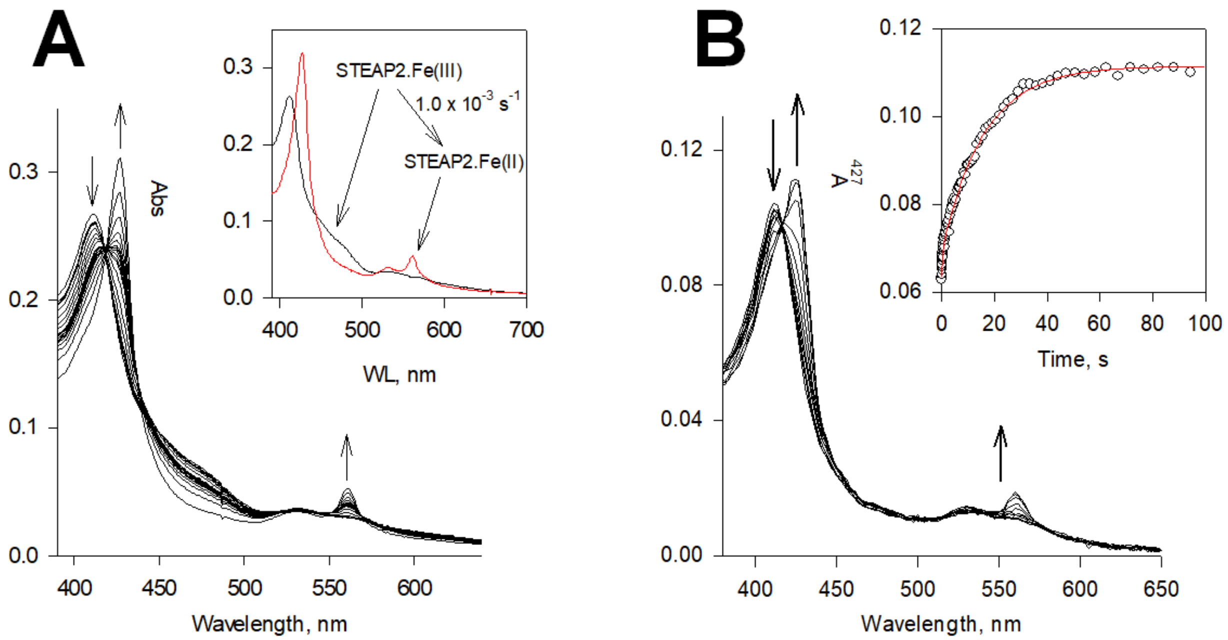


**Figure S6. Reduction of the heme on STEAP2.** (A) STEAP2, 2.3 μM, was pre-incubated with 2.5 μM FAD and reacted anaerobically with 45 μM NADPH; the spectral change was monitored for 1230 s. Inset, the resolved spectral species by deconvolution together with the conversion rate constant. Black: the ferric STEAP2 with FAD and red, the ferrous STEAP2 with reduced FAD. (B) Rapid-scan reaction of 1.1 μM STEAP1 with 4.5 μM reduced FAD; the spectral change was monitored for 100 s. Inset, the time course of $A_{427}$, the Soret absorbance of ferrous heme, extracted from the rapid-scan data. Red: biphasic exponential fit with rate constants of 2.9 (± 0.80) and 0.069 (± $3.3 \times 10^{-3}$) $s^{-1}$, respectively (n = 6). The percentage of each phase is 16% and 84%, respectively.

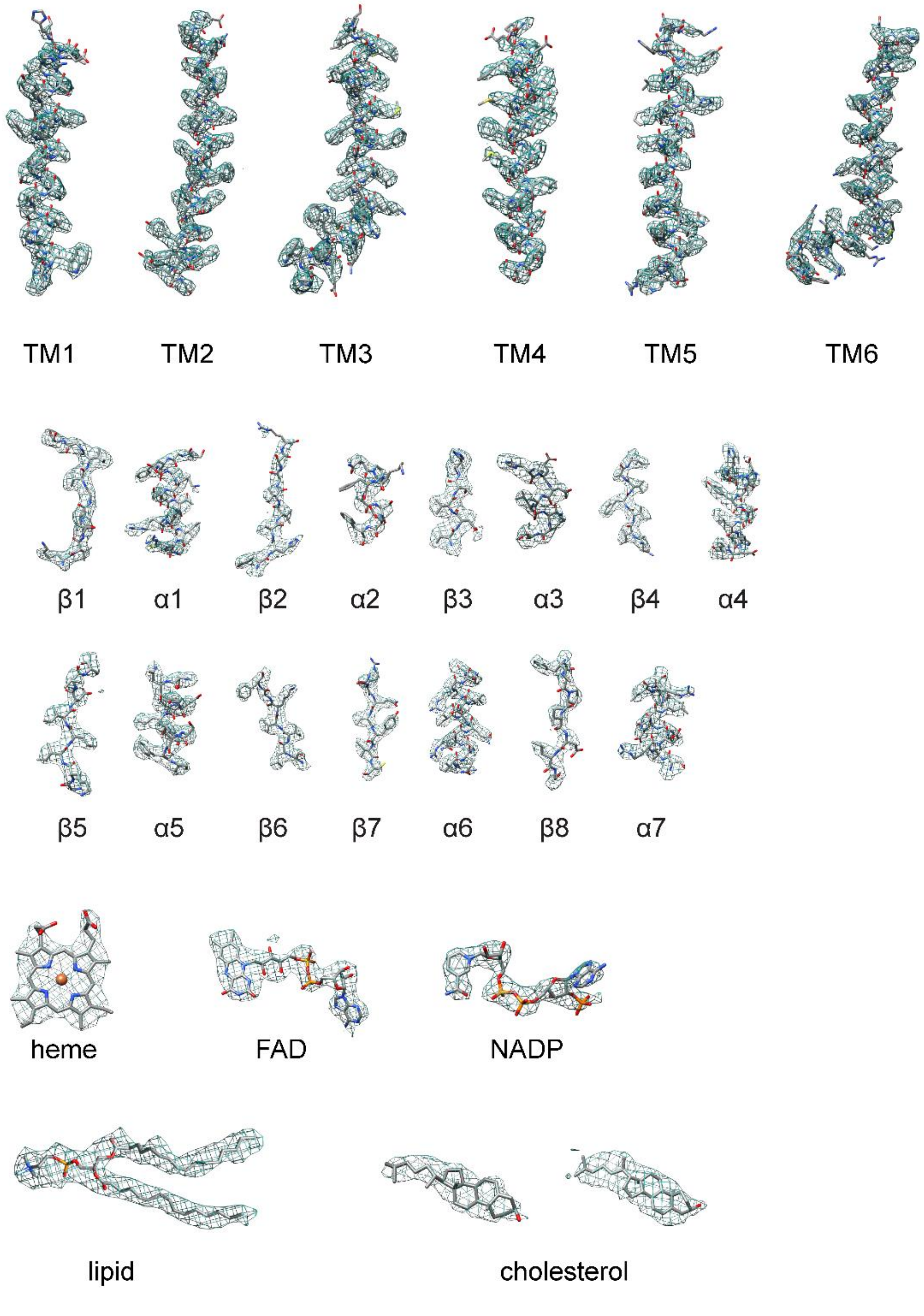


**Figure S7. Density maps of the structural elements, heme, FAD, $NADP^+$, and lipids in the cryo-EM of hSTEAP2.**

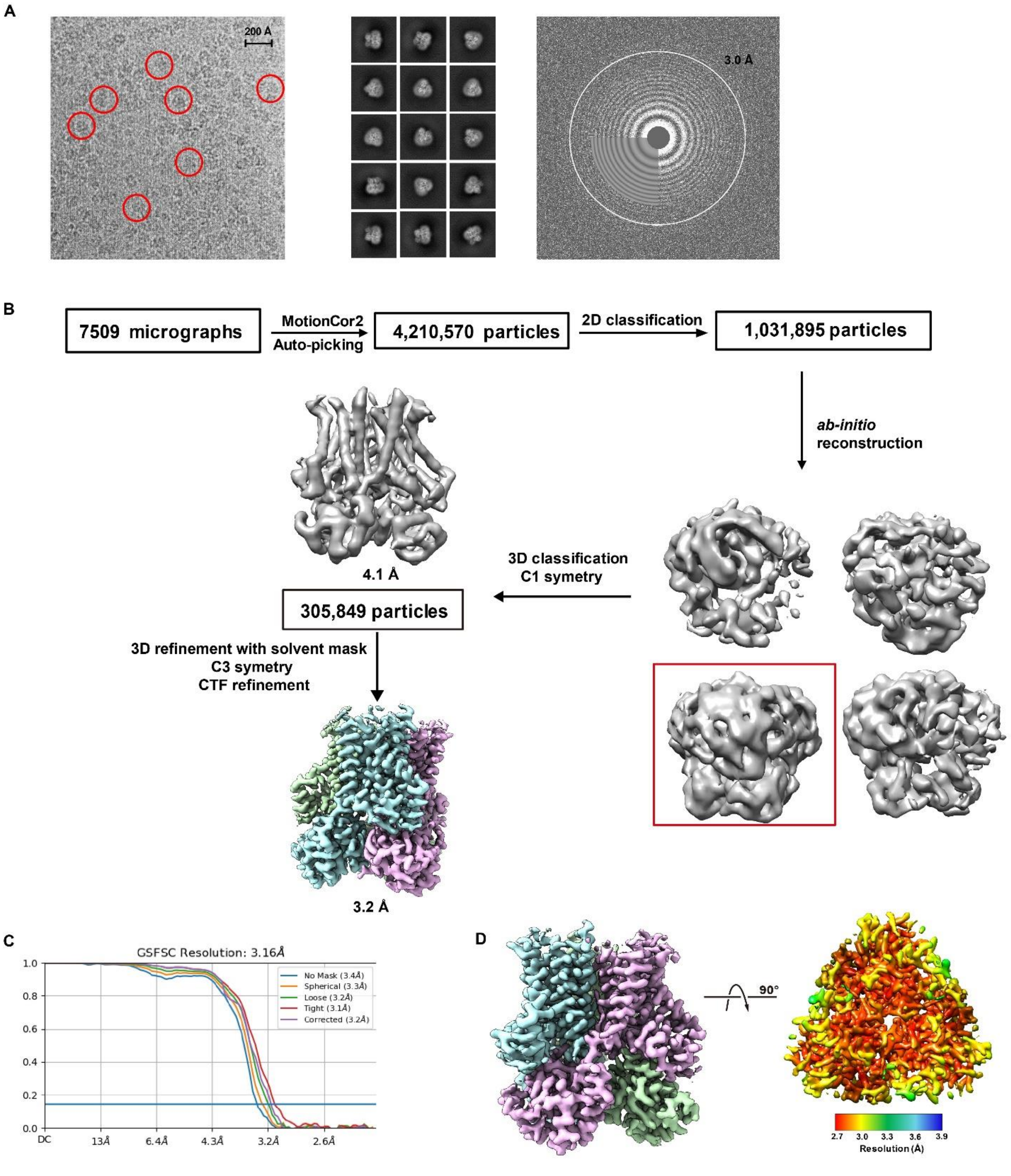


**Figure S8. The images (A) and processing (B - D) of the EM data of hSTEAP2.**
